# Protection against Poly Microbial Sepsis by Chitin oligomers is fine tuned by N-Acetyl D-Glucosamine residues

**DOI:** 10.1101/2024.10.10.617521

**Authors:** Paresh Patel, Geetanjali Agnihotri, Shilpa Siddappa, Balachandran Ravindran, Taslimarif Saiyed

## Abstract

Chitin, poly-N Acetyl D-glucosamine, is an abundant polysaccharide produced by fungal cell walls, insect epicuticle and nematodes cuticles. Its immunomodulatory function has curious attributes - paradoxically opposing host responses using different host receptors resulting in immunostimulatory or immunosuppressive propertis depending on the size/length of the oligomers as well as aceytylation levels have been reported in literature. Here we demonstrate that Hepta-N-Acetyl Chitoheptaose (7 mer) is a TLR2 ligand, 8 mer activates immune cells through TLR4 while, 6 mer is a relatively poor ligand for both TLR2 and TLR4 in generating inflammatory host cytokines. Significantly enhanced inflammatory response characterized by increased TNF-a, IL-1b, IL-6 and IL-10 was a feature of only 8 mer while, 6 and 7 mers induced modest activation in both HEK cells (transfected with specific TLRs) and in human THP2 cells in vitro. The translational significance of these features were addressed in an experimental model of Sepsis using Cecal Ligation Puncture (CLP) protocol, a murine model, considered a gold standard for human Sepsis. More significantly, therapeutic rather than prophylactic administration, that simulates real life scenerio of human and animal sepsis, of 7 mer rather than 6 or 8 residues of Chitin oligomer, significantly protected mice against sepsis as shown by decreased mortality and decreased induction of inflammatory cytokines. These findings suggest that modulation of immune response by chitopolysaccarides *in-vivo* is precisely caliberated.

## Introduction

Chitin, the second most abundant molecule that nature makes (next to Cellulose) has been a subject of intense study for their immunomodulation properties. Studies have shown that chitin can attract and activate innate immune cells leading to induction of activation of macrophages by classical pathway leading to inflammatory cytokine and chemokine production upon recognition by Pattern Recognition Receptors ^1^. It has also been shown to induce accumulation of alternatively activated macrophages, eosinophils and basophils expressing IL-4, that display anti-inflammatory properties similar to helminth and allergic immunity ^2^. The basis of this dichotomy in their biological activity reported literature awaits detailed exploration. The interaction of chitin with the immune cells and their modulation was found to be dependent on both the size of the chitin particles as well as the its source ^3,4^ where particles larger than 40 µm induce an allergic response (classical Th2) while smaller particles (1–10 µm) induce both anti-inflammatory and protective Th1 responses ^4^. Further, the source of chitin also contributes to diversity of biological activity - immunomodulation by the chitin particles derived from the crab shells varied significantly from those derived from C. albicans yeast (fungal) cells owing to the difference in their composition ^3^. Apart from length of the chitin homo-polymer, its degree of acetylation was another factor that was found to contribute to its immunomodulatory properties ^5^. Studies performed using chitosan (deacetylated version of Chitin) and the chito-oligomers reported significantly variable biological activities such as anti-microbial, anti-cancer and anti-inflammatory etc., ^6^ Degree of acetylation of Chito-oligomers of 2-6 chain length viz., 0%, 50% and 85% neutralised LPS mediated IL-6 and Nitric Oxide secretion variably, where 12% acetylated chito-oligomers showed higher reduction in the pro-inflammatory cytokines and alleviated the effects of endotoxemia in mice ^7^. The studies identified six-subunit-long chitin oligomers as the smallest immunologically active motif and that the more defined and size controlled, C10-15 chito-oligomer mixture activated myeloid, lymphoid innate and adaptive immune cells. C10-15 chito-oligomers activated Natural Killer (NK) cells and B lymphocytes,and maturation of dendritic cells enabled CD8 T cell recall responses ^5,8^. These investigations had used either cell lines or primary cells of animal or plant origin. All the contradictory observations made during the last few decades on immune-modulation by the chito-oligomers or the chito-polysaccharide have limitted the field from development of drugs and potential clincial use in human and veternary practice. The primary objectives of the current study were two fold – a) to dissect the role of specific TLRs involved in the context of the size and number of residues of chito-polysaccahrides by using HEK cells stably transfected with specific TLRs to offer clarity to controversial observations reported in literature and more critically at a translational level to b) evaluate the differential ability of chito-polysaccharides of variable residues for treatment and management of clinical sepsis in a mouse model of poly microbial sepsis viz., cecal ligation and puncture (CLP), considered to be a gold standard for pre-cinical evaluation of potential drugs against sepsis/septic shock.

## Results

### Acetylation is critical for immunomodulatory properties of chito sugars

The role of acetylation in immunomodulatory activity of chito-polysaccahride was addressed by using a preparation designated as GA120 after complete deacetylation of commercial chitosan and complete aceytylation of GA120 designated as GA132 as mentioned under methods. These two forms of chito-polysaccharede were tested in vitro using THP-1 cell line and their activation status was scored-bacterial endotoxin (LPS) was used as a positive control. The results shown in Fig 1A reveals the criticality of acetyl groups in chito-polysaccahrides in inducing the follwing cytokines/chemokines: IL-6, IL-1b, TNF-a, IL-10 and MCP-1.

**Fig 1.**
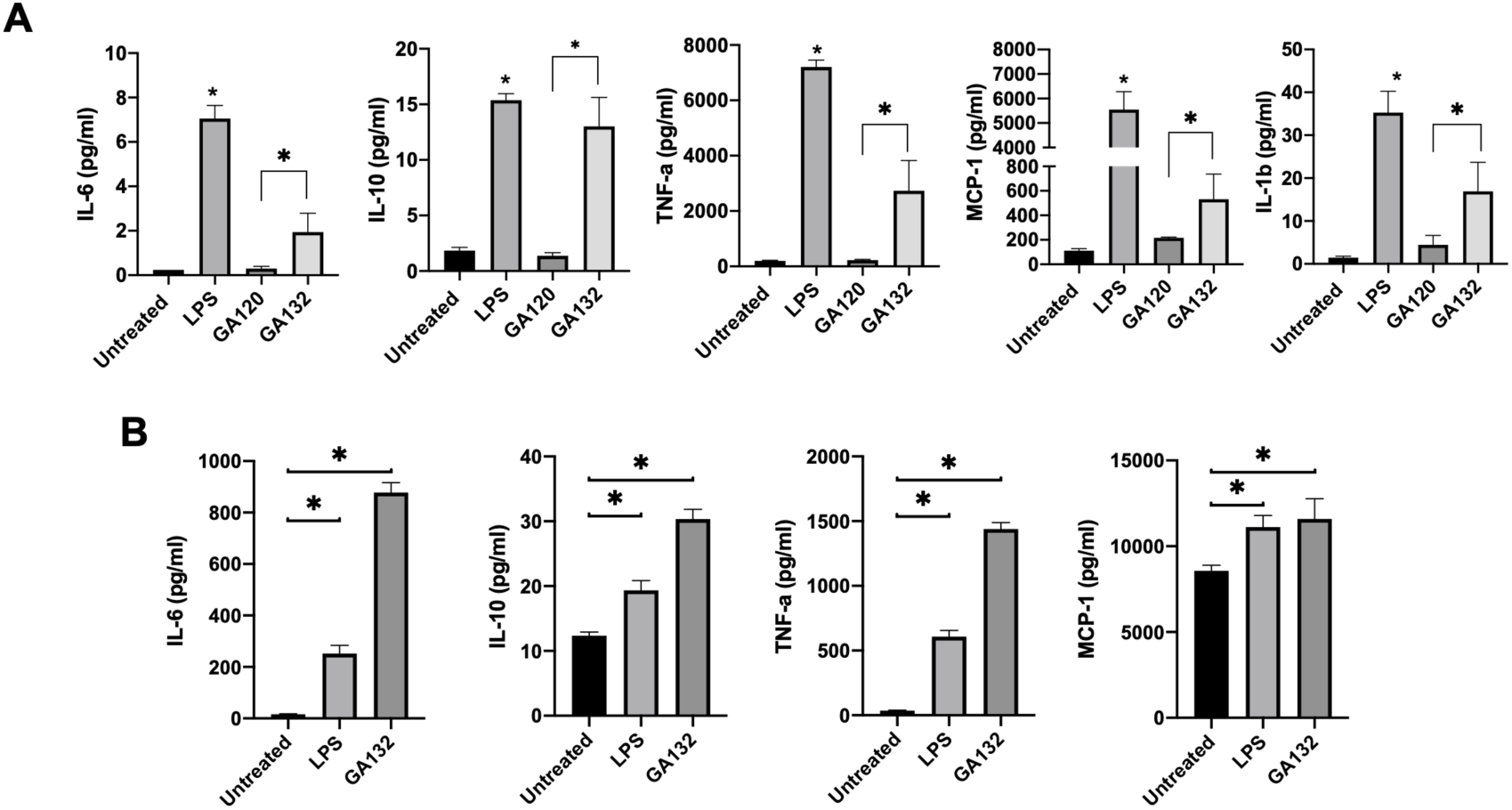
Acetylation of chito-oligomers is critical for their immunomodulatory property. **A.** Effect of GA132 (completely acetylated chito-oligomer mixture with 4-8 mer chain length) and its deacetylated version, GA120 (generated by methanolic NaOH treatment) was studied on monocytic cell line, THP-1. 1×10^6^ cells were seeded in a 24-well plate and treated with 50ug/ml of either GA132 or GA120. 100ng/ml LPS was used as a positive control. The culture supernatant from three independent biological replicates was collected and used for analysing the indicated cytokines using Bio-Plex assay. **B.** Effect of GA132 was studied on primary blood cells. 1×10^6^ PBMCs were seeded in 24 well plate and treated with 50 ug/ml of GA132 on cytokine secretion from treated PBMCs. 100ng/ml LPS was used as a positive control in the assay. The culture supernatants were collected, and levels of IL-6, IL-10, TNF-a and MCP-1 were determined using Bio-Plex assay. The data represent mean with SD from three independent biological replicates.

These observations were consistent with findings on long chain acetylated and deacetylated chito-oligomers, C10-15 and zymosan reported earlier ^5^. Since THP-1 cell lines are immortalised human monocytes the importance of acetylation was further confimed by using human primary cells (PBMCs) and the results are shown in Fig1B - the ability of GA132 in activating PBMCs was comparable or superior to the positive control, LPS in terms of potent induction of IL-6, IL-10, TNF-a and MCP-1. These results confirmed the importance of acetylation and that GA132 was indeed capable of immunomodulation of innate immune response with simultaneous secretion of both pro and anti-inflammatory cytokines and could assist in restoring the immune homeostasis in vivo in a mouse model of sepsis viz., cecal ligation and puncture ^9^.

### GA132 protects mice in cecal ligation and puncture model

The protective nature of GA132 was studied by administration of GA132 in two different strains of mice - BALB/c and C57BL/6 (Fig 2A and 2B respectively) since the two strains of mice display different levels of susceptibility in CLP model. In Balb/c mice, treatment was performed by administration 500ug GA132 along with standard antibiotic treatment (Fig 2A). A two different doses of GA132, 300ug and 700ug were administered in C57BL/6 - mice treated with higher dose showed bettwer survival (Fig 2B).

**Fig 2.**
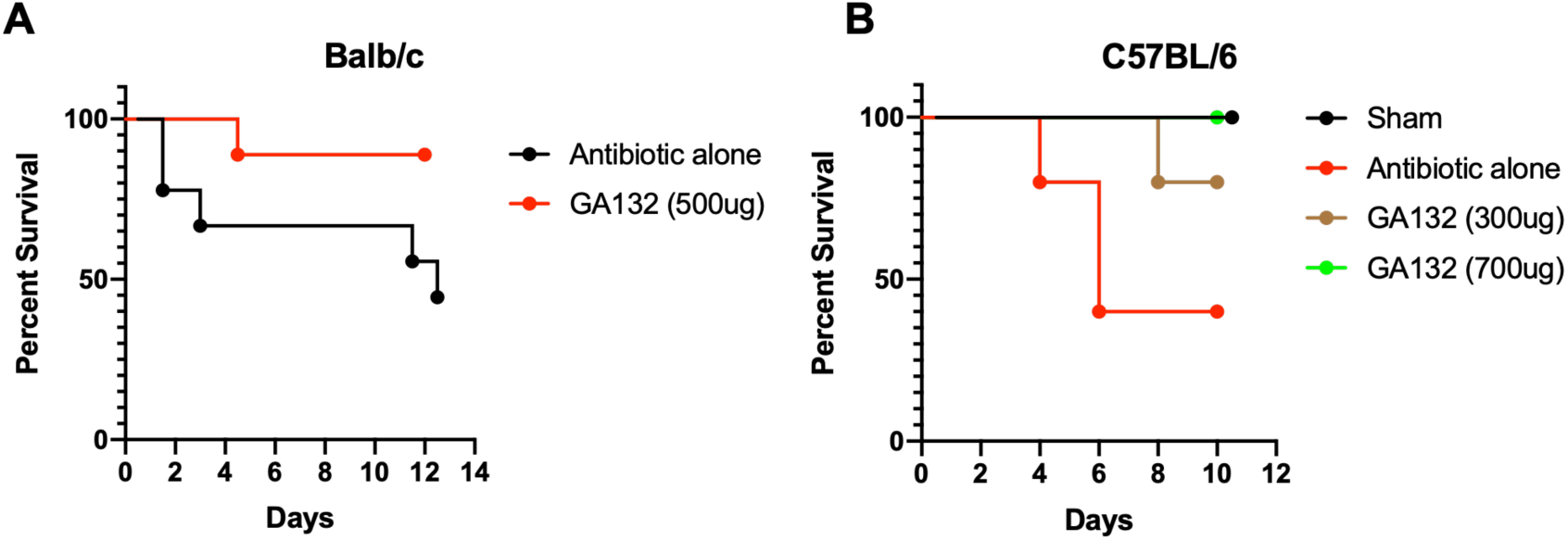
Acetylated Chito-oligomers (GA132) protects mice from sepsis in cecal ligation and puncture model. **A.** BALB/c mice (n=18) were subjected to CLP and randomly divided in two group with 9 mice in each group. CLP was followed by treatment with antibiotics along with, either 500ug of GA132 (GA132) or control PBS (antibiotic alone) and the mortality was scored over a period of 12 days. The control group showed more than 40% mortality in the first 48 hours of CLP with more than 60% overall mortality. The group treated with GA132 showed about 90% survival over 12 days. **B.** C57BL/6 mice (n= 20) were subjected to sham or CLP surgery. The CLP mice were randomly divided in 3 groups with 5 mice in each group. Surgery was followed by treatment with antibiotics along with, either GA132 (GA132 300ug or GA132 700ug) or control PBS (antibiotic alone) and the mortality was scored over a period of 12 days. The control group showed upto 60% mortality by day 6 while both the treatment groups showed protection ranging between 80-100% over the study period.

### Chito-polysaccharide composition analysis of GA132

The above observations on immunomodulation and protection mediated by GA132 suggested the need to undertake detailed molecular analysis of chito-polysaccahrides in the preparation of GA132. The MALDI-ToF analysis revealed presence of fully acetylated chito-oligomers with chain length from of 4 to 8. The major species identified were with mass of 1056.3, 1259.5, 1462.4 and smaller quantities of 1665.7 corresponding to sodium salt of Penta N-acetyl Chitopentaose (5-mer), Hexa N-acetyl Chitohexaose (6-mer), Hepta N-acetyl Chitoheptaose (7-mer) and Octa N-acetyl Chitooctaose (8-mer) respectively (Fig 3A). These results were confired by HPLC results, where GA132 showed overlapping peaks corresponding to Hexa-N-acetyl Chitohexaose (6-mer) and Hepta-N-acetyl Chitoheptaose (7-mer) along with small chain oligomers (Fig 3B).

**Fig 3.**
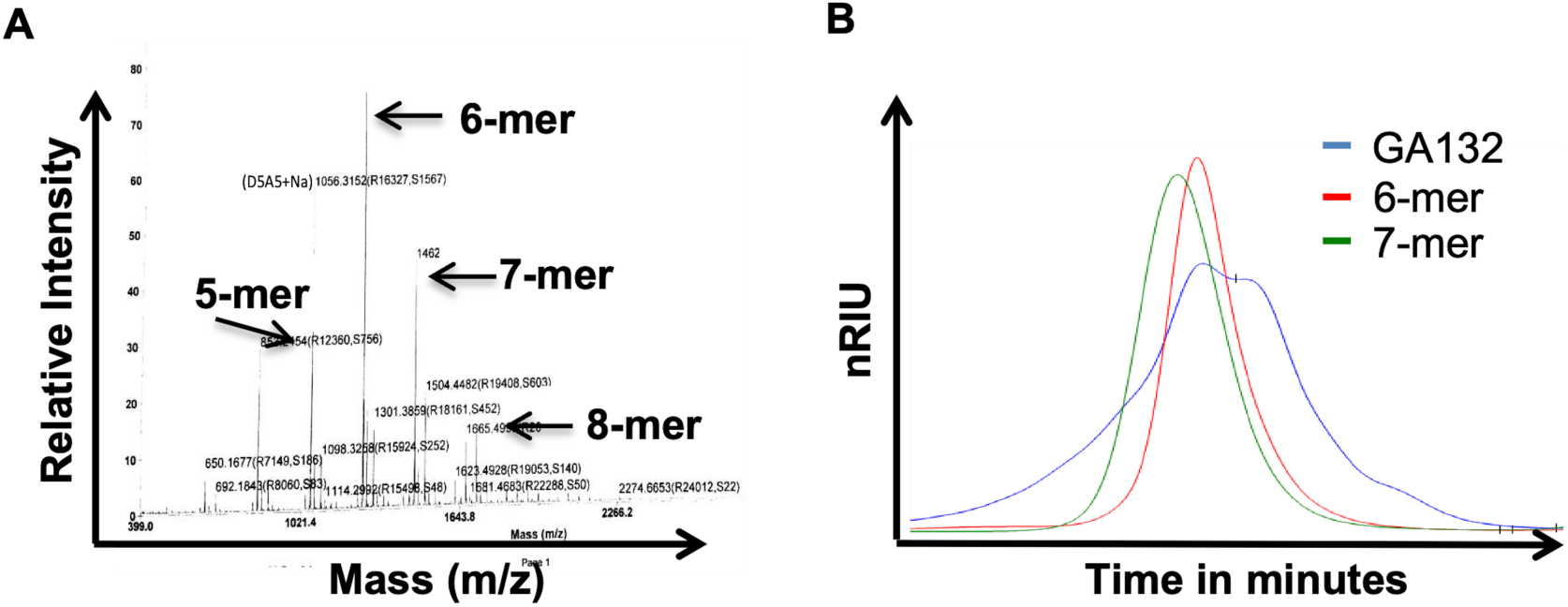
Chito-polysaccharide composition analysis of GA132. MALDI-TOF (A) and HPLC (B) analysis of GA132 indicating the presence of different chain lengths (4-8 mer) of oligomer present in the sample with 5,6 and 7-mer being dominant chain lengths

### 6-mer, 7-mer and 8-mer are qualitatively different in terms TLR usage for cellular activation

Earlier reports in literature suggested that a minimum chain length of 6 or more is required for binding and activation TLR2 ^5^ and that the chain length of the Chito-polysaccharide plays an important role in their bio-activity ^10,11^. To further delineate the activity of individual components of GA132, purified 6-mer, 7-mer and 8-mer were tested for their effect on different TLRs using specific HEK-Blue reporter cells stably transfected with TLR-2, TLR-3 and TLR-4 (Fig 4A-C) - Pam3CSK4, Poly I:C and LPS were used as positive controls respectively. Such reporter assays offer precise usage of receptors for different ligands leading to secretion of cytokines like IL-1b, IL-6, TNF-a and MCP-1, IL-10 ^12–14^. The results indicated that GA132, 7-mer and 8-mer activated TLR2, while 6-mer had no significant effect on TLR2 based NF-kB and AP-1 activation (Fig 4A). In another study, the chitohexaose derivative, AVR-25 didn’t show any binding affinity to TLR2 ^15^. Activation thro TLR2 was comparable to GA132 only by 7-mer while 6 mer failed to activated TLR2 transfected cells. GA132, and 8- mer activated TLR-4 transfected HEK-cells with poor activation by 6-mer while 7-mer had not no activity (Fig 4C).

**Fig 4.**
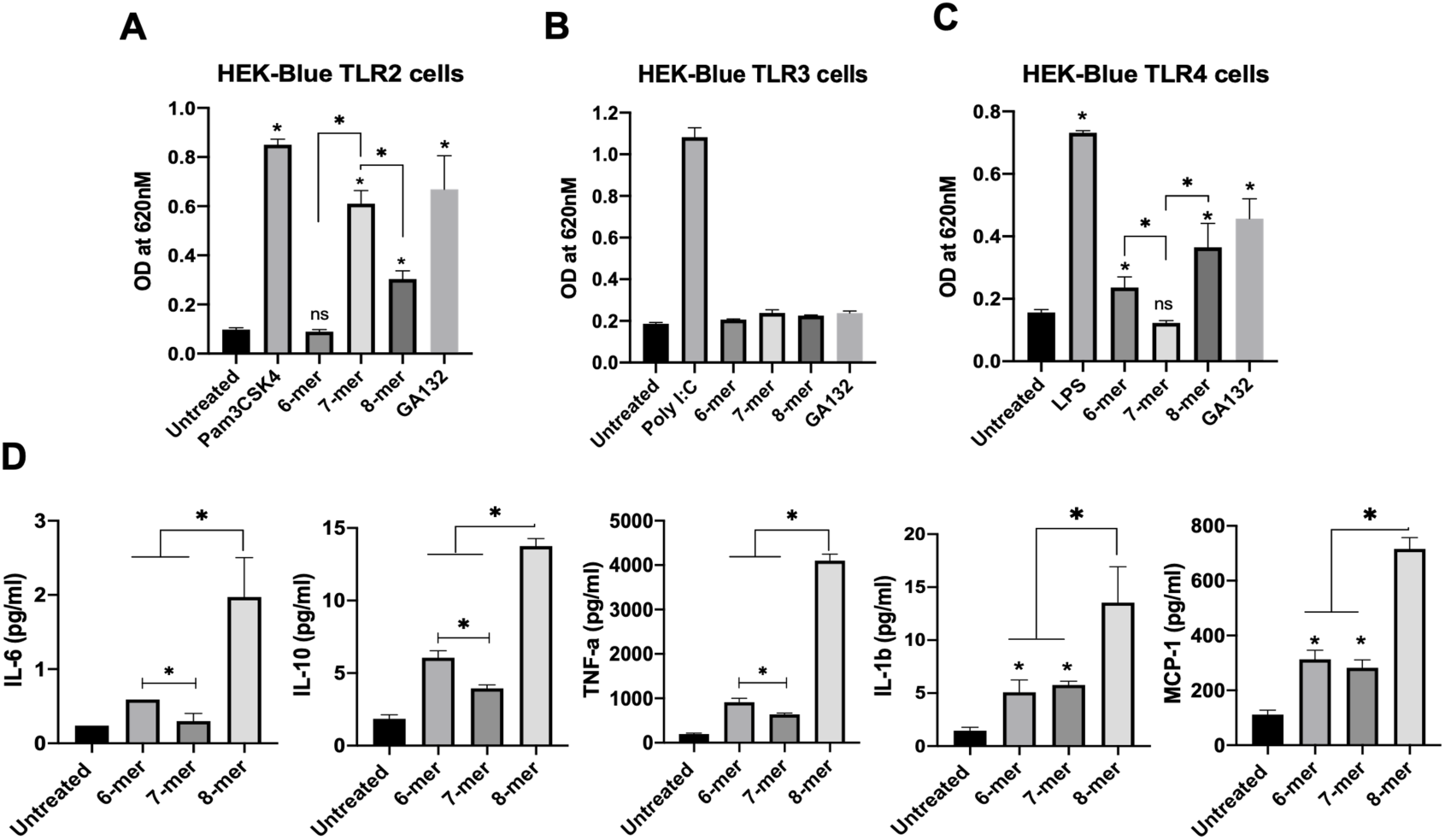
Purified N-acetyl chito-polysaccharide of different chain length are qualitatively different in terms of TLR usage for cellular activation. **A,B and C.** The effect of GA132 and purified 6-mer, 7-mer and 8-mer were tested on different TLRs was studied using HEK-Blue TLR reporter cells. 2×10^4^ HEK-Blue TLR2 (A), TLR3 (B) and TLR4 (C) reporter cells were seeded in 96 well plates and the treated with 50ug/ml of sample and absorbance at 620nm was measured 16 hr post stimulation. The data represents Mean with SD from biological triplicates from one of the multiple experimental sets. PAM3CSK, Poly:IC and LPS were used as positive controls in TLR2, TLR3 and TLR4 assays respectively. **D.** Effect of 6-mer, 7-mer and 8-mer purified oligomers was studied on THP-1 cells. 1×10^6^ cells were seeded in a 24-well plate and treated with 100ug/ml of oligomers. The culture supernatants were collected, and levels of IL-6, IL-10, TNF-a and MCP-1 were determined using Bio-Plex assay. The data represent mean with SD from three independent biological replicates.

None of the chito-oligomers or GA132 activated HEK cells transfected with TLR3 (Fig 4B). To further explore the effect of this differential activation of TLRs by 6-mer, 7-mer and 8-mer, THP-1 cells were treated with purified oligomers and the cytokines levels were quantified after 24 hours (Fig 4D). The 6-mer and 8-mer induced significantly higher levels of IL-6, IL-10 and TNF-a than 7-mer. 8-mer treatment showed the highest induction of all the tested cytokines/chemokines (Fig 4D). These results confirmed that chain length chito-oligomers were qualitatively variable in terms of receptor usage, cellular activation and release of inflammatory cytokines.

### In vivo protection against murine sepsis by Hepta N-acetyl Chitoheptaose (7-mer) is superior to 6 mer and 8 mer

Since Chitin oligomers have the potential to modulate immune responses of the host their utility for modifying diseases such as Sepsis have been studied by several investigators ^7,10,16^ LPS mediated endotoxemia model in mice has been widely used nothwithsatnding the limittation that the model does not truely represent sepsis. In this study protective effects of purified 6-mer, 7-mer and 8-mer were assessed in the cecal ligation and puncture (CLP) model in mice, considered to be a gold standard as pre-clinical model of sepsis. Intially the therapeutic efficay was tested in groups of mice by administration of defined chito-oligosaccharides 6 hrs post CLP procedure and the survival in different groups was evaluated over a period of 7 days,. The 7-mer protected 90% of mice from mortality as shown in Fig 5A and protection induced by 6-mer and 8-mer was not significant in comparison to controls. Chitohexaose (6-mer) has been reported earlier to induce significanct protection in a model of endotoxemia in which purified LPS was used to challenge mice^10^. The oberved difference between the two studies could be due to use of different model systems - LPS induced endotoxemia versus CLP. The latter model more closely simulates polymicrobial sepsis in which multiple pathways are activated ^17^. In another study, a derivative of chitohexaose was found to protect young and aged mice in the CLP model ^15^, in which the underivatized chitohexaose was not tested. The therapeutic efficay of 7-mer could be demonstrated even if the treatment was performed 24 hrs post CLP procedure as shown in Fig 5B. About 80% survided when treated with 7-mer in comparison to control groups while 6-mer and 8-mer were far less effective in mediating protection.

**Fig 5.**
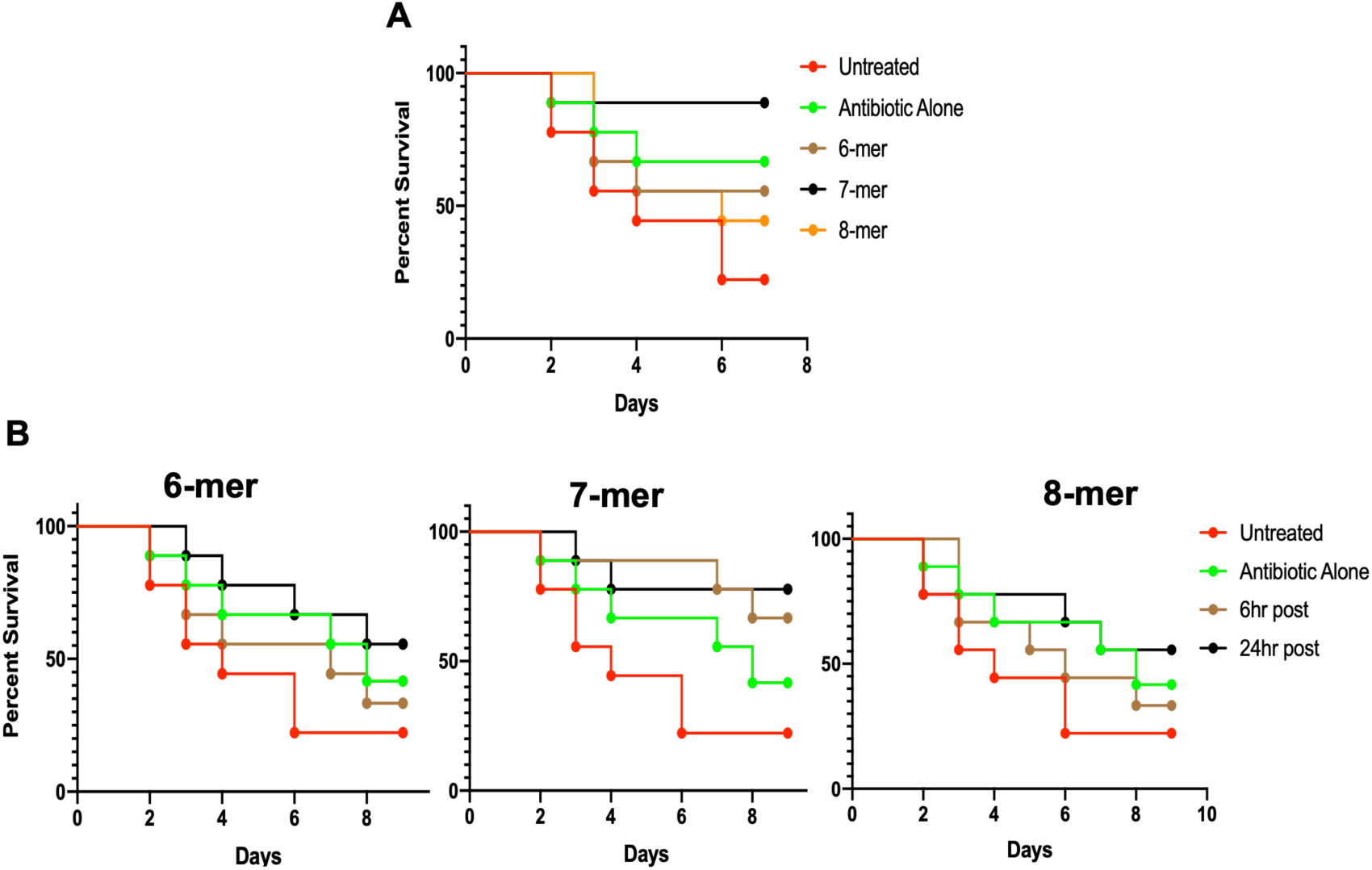
Hepta N-acetyl Chitoheptaose (7-mer) provides superior in-vivo protection against murine sepsis. **A.** C57BL/6 mice (n= 45) were subjected to sham or CLP surgery. The CLP mice were randomly divided in 5 groups with 9 mice in each group. The untreated group received no antibiotics and 6 hours post-surgery, the other groups were treated with antibiotics along with, either single dose of 250ug of oligomers (purified 6-mer, 7-mer and 8-mer) or control PBS (antibiotic alone). The mortality was scored over a period of 7 days. The untreated group showed 80% mortality over 7 days, while in the 7-mer treated group 8 out of 9 mice survived. **B.** Comparison of effect of purified 6-mer, 7-mer and 8-mer intervention at early (6 hours) vs late (24 hours) time points post CLP in C57BL/6 mice (n=9 for each group). The three additional groups were added to the CLP study groups used in 3A, where the mice received in intervention with single dose of 250ug of oligomers (purified 6-mer, 7-mer and 8-mer) 24 hours post CLP. The mortality was studies over 9 days and data was compared with the different time of intervention with the same molecule.

### Hepta N-acetyl Chitoheptaose (7-mer) reduces cytokine storm

Recent Clinical studies have indicated that a cytokine network of IL-6, IL-8, MCP-1 and IL-10 plays a pivotal role in the acute phase of sepsis and higher levels of IL-6, TNF-a, IL1-b and MCP-1 were associated with higher mortality in sepsis patients^18,19^. Treatments that reduced induction of inflammatory cytokines significantly improved the survival in mice models ^10,15,20^. Hence, plasma levels of these cytokines at 6 and 48 hours post treatment were analyzed from CLP mice treated with 6-mer, 7-mer and 8-mer where, the treatment was administered at 6 hours post CLP and plasma levels of cytokines were tested 48hr post CLP. The results revealed consistant decrease of IL-6, IL-1b, TNF-a and MCP-1 in comparison to control mice (Fig 6A). Analysis of IL-10, TNF-a, IL-1b and MCP-1 at 48 hours post surgery showed significant reduction only in 7-mer treated group when compared to untreated groups (Fig 6A).

**Fig 6.**
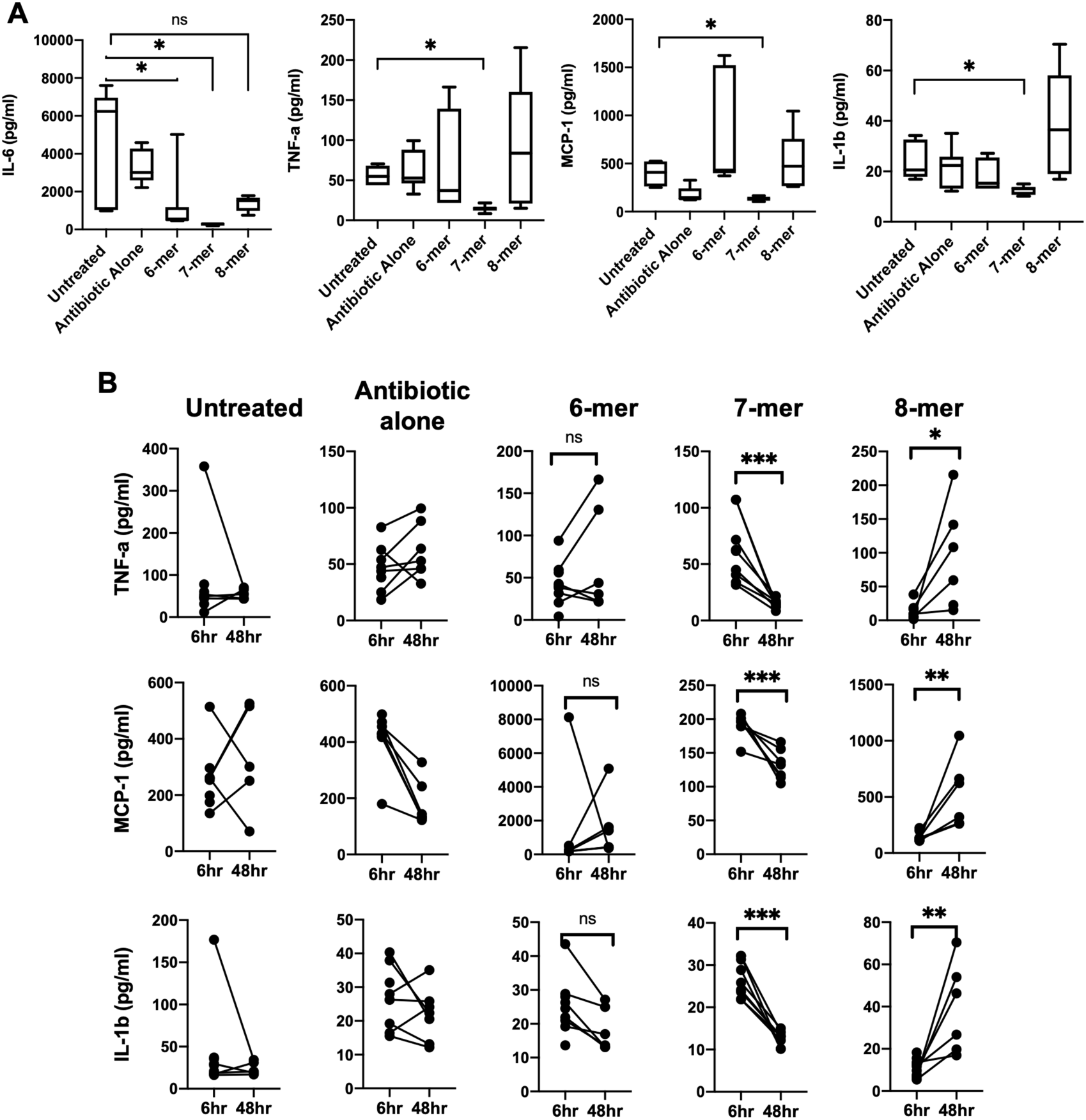
Hepta N-acetyl Chitoheptaose (7-mer) reduces cytokine storm. **A.** Plasma was collected at 6 hours and 48 hours post CLP from different treatment groups in 5A. The plasma was analysed for IL-6 from 6 hours samples (n=9 in each group) and from 48 hours samples; levels of TNF-a, IL-1b and MCP-1 were determined using Bio-Plex based assays. n=5 for untreated group, n=6 for antibiotic alone, n=6 for 6-mer group, n=8 for 7-mer and n=6 for 8-mer group. **B.** Effect of 6-mer, 7-mer and 8-mer on progression of TNF-a, MCP-1 and IL-1b over time in individual mice. Plasma was collected at 6 hours and 48 hours post CLP from different treatment groups in 3A. n=9 in each group at 6 hours and for 48 hours samples; n=5 for untreated group, n=6 for antibiotic alone, n=6 for 6-mer group, n=8 for 7-mer and n=6 for 8-mer group.

Finally, effect of 6-mer, 7-mer and 8-mer on progression of inflammatory cytokines were compared in plasma samples collected at two time points post CLP susgery viz., 6hrs and 48 hrs. The chito-oligosaccahrides were administered 6 hrs post surgery. The results shown in Fig 6B revealed significant reduction in levels of TNF-a, IL-1b and MCP-1 over time in only 7- mer treated group, while 8-mer treatment led to increase in the levels of these cytokines in the same period. 6-mer treatment had no effect on the levels on these cytokines (Fig 6B). The sustained increase in pro-inflammatory cytokines correlate well with differences observed in survival of mice shown in Fig 5A. These in vivo findings in CLP model of mice are consistent with the in-vitro released cytokine levels in THP-1 cells by chito-oligosaccahrides of 6,7 and 8 residues (Fig 4D).

## Discussion

Chitosan, a largely deacetylated form of chitin has been tested widely as an immunomodulator ^16,21^ while in the current study aceytylated residues have been tested after establishing the importance of aceytylation. Significant protection was mediated in a pre-clinical model of Sepsis in mice by 7-mer chito-polysaccharide and not by 6-mer or 8-mers. Similar observations were made recently in another disease model where, the effect of different chain chito-oligomers was studied in Myocarditis Model and 7-mer showed better activity in-vivo than 6-mer and 8-mer ^11^.

Differences in absorption and bio-availability of chito-oligomers based on size along with capability to stimulate different TLRs have been reported ^11,22^- the observations in the current study clarifies the fine difference between 6-mer, 7-mer and 8-mer in specific TLR usage by them both *in-vitro* and *in-vivo*. Sepsis is a complex disease initiated by microbial infections potentiated by host inflammatory responses. Immune dysregulation leads to organ failure leading to mortality ^23–25^. While microbial infections are generally controlled by use of antibiotics, bad prognosis leading to mortality is largely attributed to immune-paralysis and immunotherapy to modulate the host immune response ^26,27^. TLR4 has been explored widely as a potential target in sepsis and various molecules, which antagonize and restrict TLR4 activation, for sepsis intervention ^28^. Although, some of these LPS antagonists were promising in pre-clinical studies, clinical trials failed consistently ^29^. TLR2 is another receptor that plays a role in polymicrobial sepsis by regulating neutrophil migration ^30^. Compared to use of TLR-4 antagonists, immunomodulation through TLR2 to restore immunological equilibrium as a therapeutic strategy for sepsis has not been explored more extensively. Prophylactic treatment of chitosan oligosaccharides were found to attenuate inflammation and oxidative stress and protect mice from LPS challenge ^16^. Chitohexose has been demonstrated to block endotoxemia by induction of alternate activation of macrophages etc^10^. The deacetylated or partially acetylated chito-oligomers possess different anti-oxidant and anti-inflammatory activities based on mitogen-activated protein kinase (MAPK) signaling^31^. Unfortunately most of the biological studies on immunomodulation have been performed with ill-defined chitin oligomers with variable levels of acetylation. Taken together, the following aspects stand out as novel and definitive findings of the current study - 1. Use of different TLR usage by 6-mer, 7-mer and 8-mer of chito-polysaccharides in activating immune cells, thus demonstrating precise ligand and receptor specificity; 2) Criticality of acetylation and number of residues of N-Acetyl D-Glucosamine in chito-oligomers modulating variably inflammatory cytokines; 3) Induction of significant *in-vivo* protection by 7-mer in a CLP model of sepsis and; 4) Protection against murine sepsis by therapeutic administration of 7-mer (as opposed to prophylactic administration performed widely by investigators), simulating real-life scenerio in human sepsis when patients report after onset of clinical sepsis. The mechanism by which 7-mer residue of Chito-polysaccahride recaliberates immune tolerence in sepsis needs to be studied in a manner similar to another polysaccharide, b-glucan reversing epigenetic state of LPS induced immunological tolerance^26^.

## Materials and methods

### Cell lines and Primary cells

HEK-Blue TLR2, HEK-Blue TLR3 and HEK-Blue TLR4 reporter cells expressing SEAP enzyme under NF-kb and AP-1 promoter (Cat. no.# hkb-htlr2, hkb-htlr3 and hkb-htlr4) were purchased from InvivoGen (USA). Human Peripheral blood mononuclear cells (PBMCs) (Cat. no.# CL003-25) were purchased from Himedia laboratories (India). HEK based cells and PBMCs were cultured in DMEM and RPMI media respectively. The cells were maintained at 37^0^C incubator with 5% CO2. The culture media was supplemented with 10% Fetal Calf Serum (Gibco) along with standard antibiotics. HEK-Blue selection antibiotics (InvivoGen, Cat. no.# hb-sel, ant-zn and ant-bl) were added as per manufacturers instructions.

### Chemicals and reagents

Hexa-N-acetyl Chitohexaose (Cat. no.# 56/11-0050), Hepta-N-acetyl Chitoheptaose (Cat. no.# 57/11-0010) and Octa-N-acetyl Chitooctaose (Cat. no.# 57/12-0010) termed 6-mer, 7-mer and 8-mer respectively were purchased from Isosep, Sweden. All other chemicals were purchased from Sigma. Chito-oligomer mixtures were generated by previously used protocol using acid hydrolysis with modifications ^32^. Briefly, chitosan (Sigma-aldrich cat no. 448877) was hydrolysed with conc. HCl by heating at 72°C for 45 minutes. Same amount of water was added to the reaction mixture and kept at -20°C for two day. The precipitate was washed with chilled ethanol and acetone followed by drying under vacuum. The dried precipitate was acetylated using Ac_2_O in presence of triethyl amine. To deacetylate the hydroxyl groups, the reaction mixture was treated with methanolic NaOH. The mass profiling of the sample suggested the presence of 3-10 mers of N-acetyl glucoseamine. Different fractions were generated on Biogel - P4 (Bio-Rad and Cat. No. 1504124) and tested on reporter cells.

### TLR reporter assays

2×10^4^ HEK-Blue reporter cells were seeded, in each well of a 96 well plate, in HEK-Blue detection media (InvivoGen, Cat. no.# hb-det3) along with the 50ug/ml of the chito-oligomers. 10ng/ml Pam3CSK4 (InvivoGen, Cat. no.# tlrl-pms.), 100ng/ml Poly I:C (InvivoGen, Cat. no.# tlrl-pic.), and 10ng/ml LPS-EK (InVivoGen, Cat. no.# tlrl-eklps ), were used as a positive control for HEK-Blue TLR2, TLR3 and TLR4 cells respectively. The treated cells were incubated at 37^0^C for 16 hours and the absorbance (O.D.) was measured at 620nm in either TECAN Infinite® 200 PRO or Thermo Scientific Varioskan LUX multimode reader.

### Mice and Cecal ligation and puncture (CLP) study

The Cecal ligation and puncture (CLP) study with 7-9 week old C57BL/6 or BALB/c males was conducted, at animal house facility at TheraIndx Lifesciences Pvt. Ltd. (Bangalore, India) and Institute of Life Sciences (Bhubaneswar, India) respectively, in accordance with ethical practices laid down in the CPCSEA guidelines for animal care and use ^33^. The Institutional Animals Ethics Committee (IAEC) of the test facility approved the studies. The CLP was performed as described before ^9,34^. Sham control mice had undergone surgery with cecal manipulations without ligation and puncture. 500ul saline was administered subcutaneously immediately after surgery. Tramadol (20 mg/kg) was injected subcutaneously for post-operative analgesia. BALB/c mice were injected with single IP dose 500ug GA132 while C57BL/6 mice received single IP dose of 300ug or 700ug GA132 6 hours post CLP. A separate group of C57BL/6 mice received single IP dose of 250ug of 6-mer, 7-mer or 8-mer either 6 hour or 24 hour post CLP. All group except the CLP alone received a single dose of standard antibiotics (amoxicillin and clavulanate) 6 hour post CLP. Control group was injected with saline solution. Total of 92 C57BL/6 and 18 BALB/c mice were used in the study.

### Bio-Plex Multiplex assays

The Mutiplex immunoassay for Human and Mouse IL-1b, IL-6, IL-10, TNF-a and MCP-1 were purchased from Bio-Rad. For human cytokines, 1×10^6^ PBMCs were seeded in 24 well plate a day before and treated with 100ng/ml LPS or 50ug/ml GA132 for 24 hours. The culture supernatant from three independent biological replicates was collected and used for the assay without dilution. For mice cytokines, the plasma samples from mice were collected 6 or 48 hours post treatment and the diluted 1:2 before assay. The assay was performed as per the manufacturers protocol and the beads were read and analyzed in Bio-Plex^®^ MAGPIX™ Multiplex Reader (Bio-Rad).

### HPLC analysis

The samples were prepared in Ammonium Aceteate buffer (0.2M). 10ug of GA132 or purified standards of Hexa-N-acetyl Chitohexaose (Cat. no.# 56/11-0050), Hepta-N-acetyl Chitoheptaose (Cat. no.# 57/11-0010) were analyzed with TSK-Gel (Cat. no.# G2000SWx.) on HPLC (Agilent Technologies 1260 Infinity). The change in refractive index was observed over time for different samples. The data was plotted using (OpenLAB Control).

### Maldi-TOF

For sample analysis, 1.5 µl of 10-mg/mL α-CHCA (α-cyano-4-hydroxycinnamic acid) matrix solution was mixed with 1.5 µl of each sample. From the resulting solution, 1 µl was spotted onto the 384Opti-TOF 123mmX81mm SS target plate (AB Sciex). After drying at room temperature, spotted samples were analysed using a ABSCIEX TOF/TOF 5800 mass spectrometer (Applied Bio systems , USA) equipped with a 200 Hz, 355 nm Nd: YAG and acquired MS spectra followed by External mass calibrations for reflector mode were performed using a calibration mix (Cal mix TOF/TOF, Sciex) containing des-Arg1-bradykinin (m/z 904.468), Angiotensin I (m/z 1296.685), Glu1-fibrinopeptide B (m/z 1570.677), ACTH clip 1–17 (m/z 2093.086), ACTH clip 18–39 (m/z 2465.199), and ACTH clip 7–38 (m/z 3657.929) diluted to the manufacturers specifications (1–3 pmoL/μL each). The identified masses are exported as peek list.

### Statistical analysis

Experimental data were analyzed using GraphPad Prism 8 (GraphPad Software, Inc.). Kruskal-Wallis test and One way ANOVA test was used for analysis of cytokines and reporter assay values. Log-rank (Mantel-Cox) test was utilized for survival analysis. P-values, P < 0.05 were considered statistically significant and are denoted by * throughout if calculated P-values were lower than P = 0.05.

## Acknowlegements

The work was supported by BIRAC CRS grant (BT/CRS0135/CRS-06/14) provided to TS and BR Institute of Life Sciences is funded by core grants from Department of Biotechnology, Government of India.

## Author contributions

Conceptualization: TS,BR and PP

Methodology: PP, SS and GA

Investigation: PP, SS and GA

Visualization: PP

Funding acquisition: TS, BR and PP

Project administration: TS and RB

Supervision: TS and BR

Writing – original draft: PP

Writing – review & editing: TS, BR and PP

